# Facilitation between plants shapes pollination networks

**DOI:** 10.1101/161034

**Authors:** Gianalberto Losapio, Miguel A. Fortuna, Jordi Bascompte, Bernhard Schmid, Richard Michalet, Rainer Neumeyer, Leopoldo Castro, Pierfilippo Cerretti, Christoph Germann, Jean-Paul Haenni, Seraina Klopfstein, Javier Ortiz, Adrian C. Pont, Pascal Rousse, Jürg Schmid, Daniele Sommaggio, Christian Schöb

## Abstract

Although it is known that plant–plant and plant–pollinator interactions can strongly influence biodiversity and its effects on ecosystem functioning, the details of how competition and facilitation among plants scale up to mutualistic interactions with pollinators and thus affect pollination networks are poorly understood. We introduce a simple experimental system in which we control local plant interactions, measure pollinator responses and characterize plant–pollinator networks. We find that facilitation among plants produces synergistic and antagonistic effects on the pollinator community affecting the architecture and robustness of plant–pollinator networks. Our results provide evidence for bottom-up non-additive effects of plant interactions on pollination networks and have implications for the way we study and manage ecosystems.

---

Plants cluster together and interact among themselves and with other organisms, with fundamental consequences for biodiversity and ecological networks. However, linkages between interacting plants and plants interacting with mutualists are poorly understood in real-world ecosystems. Here, we report results of a field removal experiment with natural plant communities where we compared networks of pollinators interacting with dominant plant species, here referred to as foundation species, and their associated beneficiary plant species growing in clusters, with networks of pollinators interacting with the same foundation and beneficiary species growing alone. We tested the hypothesis that the plant–pollinator networks in multispecies clusters of foundation and beneficiary plant species are more nested and robust than the sum of networks in plant clusters containing foundation species or beneficiary species growing separately. We found that pollinator diversity and plant–pollinator network architecture were significantly different when foundation and beneficiary species grew together than what would be expected from additive effects of foundation species and beneficiary species. The directionality of these effects differed between the two foundation species used as models. Moreover, the resulting changes in network-level interaction diversity, independent from species diversity, affected simulated network robustness, with differences among extinction scenarios and foundation species. This study, therefore, sheds new light on the mechanisms behind the propagation of ecological interactions within trophic levels to interactions among trophic levels in real-world ecosystems and suggests that non-additive effects could emerge in a variety of networks of organisms and ecosystems.

Despite wide-ranging implications for biodiversity (Valiente-Banuet *et al.*, 2006; Callaway, 2007), ecosystem functioning (Hector *et al.*, 1999) and services (Schöb *et al.*, 2015; Duchene *et al.*, 2017), fundamental questions remain about the basic ecological role plant–plant interactions play in real-world ecosystems (McIntire & Fajardo, 2014). Net interactions among plant species can be positive (i.e., facilitation), neutral, or negative (i.e., competition) depending on whether the presence of interspecific neighbours enhances or diminishes the growth, survival, or reproduction of neighbours, respectively (Callaway, 2007). Independent of the underlying mechanisms, facilitation and competition mainly result in spatial aggregation (i.e., clustering) and segregation (i.e., exclusion), respectively (MacArthur & Levins, 1967; Bruno *et al.*, 2003; Meron, 2012). Facilitation is often due to the effect of dominant plant species, referred to as foundation species, which are tolerant to stress and buffer limiting environmental factors in a way that some other, subdominant associated species can benefit from the newly created environmental conditions (Bruno *et al.*, 2003; Ellison *et al.*, 2005; Callaway, 2007; McIntire & Fajardo, 2014). Generally, facilitation is now recognized as a fundamental ecological process in plant communities (Callaway, 2007) and ecosystems (Duchene *et al.*, 2017). In particular, foundation species can structure plant communities (Schöb *et al.*, 2012) by enabling species coexistence (McIntire & Fajardo, 2014) and increasing plant diversity (Hacker & Gaines, 1997; Michalet *et al.*, 2006; Cavieres *et al.*, 2014).

Few previous studies have investigated the impact of interactions between plants on pollinators (Feldman *et al.*, 2004; Molina-Montenegro *et al.*, 2008; Sieber *et al.*, 2011; Ruttan *et al.*, 2016; Mesgaran *et al.*, 2017), highlighting the linkages between the structure of plant and insect communities. However, no research has experimentally examined how facilitative plant–plant interactions may propagate to other trophic levels and shape e.g. plant pollinator network architecture.

The growing interest for positive plant interactions coincides with the growing evidence about the role mutualistic interactions play in biodiversity maintenance (Bastolla *et al.*, 2009; Bascompte & Jordano, 2014). Indeed, simultaneously but independently, the study of mutualistic networks among plants and animals has illustrated ecological and evolutionary processes shaping communities and ecosystems (Bascompte & Jordano, 2014). Differences between those two fields reside in that plant ecology has hardly considered interaction networks within plant communities (but see e.g. Losapio & Schöb, 2017) while studies applying interaction networks mainly focused on interactions between trophic levels (but see e.g. Verdu & Valiente-Banuet, 2008). Consequently, there is a lack of studies experimentally investigating how interactions among plants scale up to plant–animal networks in real-world ecosystems. In particular, we do not know to what extent plant facilitation has bottom-up cascading effects shaping the architecture and robustness of mutualistic plant–pollinator networks.

To this end, we conducted a field removal experiment with two foundation species (*sensu* Ellison *et al*., 2005) (*Arenaria tetraquetra* spp. *amabilis*, hereafter *Arenaria* and *Hormathophylla spinosa*, hereafter *Hormathophylla*) and eight beneficiary species (Fig. SI1) in the Sierra Nevada Mountains (Spain), where the importance of plant facilitation for community structure is well documented (Schöb et *al.*, 2012, 2013a,b, 2014). We assembled plant communities with foundation species and beneficiary species growing together in clusters (i.e., simulating facilitation effects) and the same foundation species and beneficiary species growing separately (i.e., simulating the two different parts of the facilitative system in isolation) and we recorded plant–pollinator interactions. To experimentally test the hypothesis that pollination networks of facilitation-driven plant clusters are more diverse, nested and robust than what would be expected from summing pollination networks of foundation and beneficiary species growing separately (Figure 1), we compared the observed network in multispecies plant clusters con-taining foundation and beneficiary species growing together (i.e., the ‘facilitation’ treatment) with the expected pollination network calculated as the sum of those found on plant clusters containing foundation or beneficiary species growing separately (hereafter referred to as ‘additive’ treatment). Finally, to test whether differences between the expected additive network and the observed network in plant clusters with foundation and beneficiary species were due to the establishment of new interactions, i.e., interaction rewiring, rather than species turnover, we considered the pollination networks composed only by pollinator species common to both treatments.

**Figure 1:**
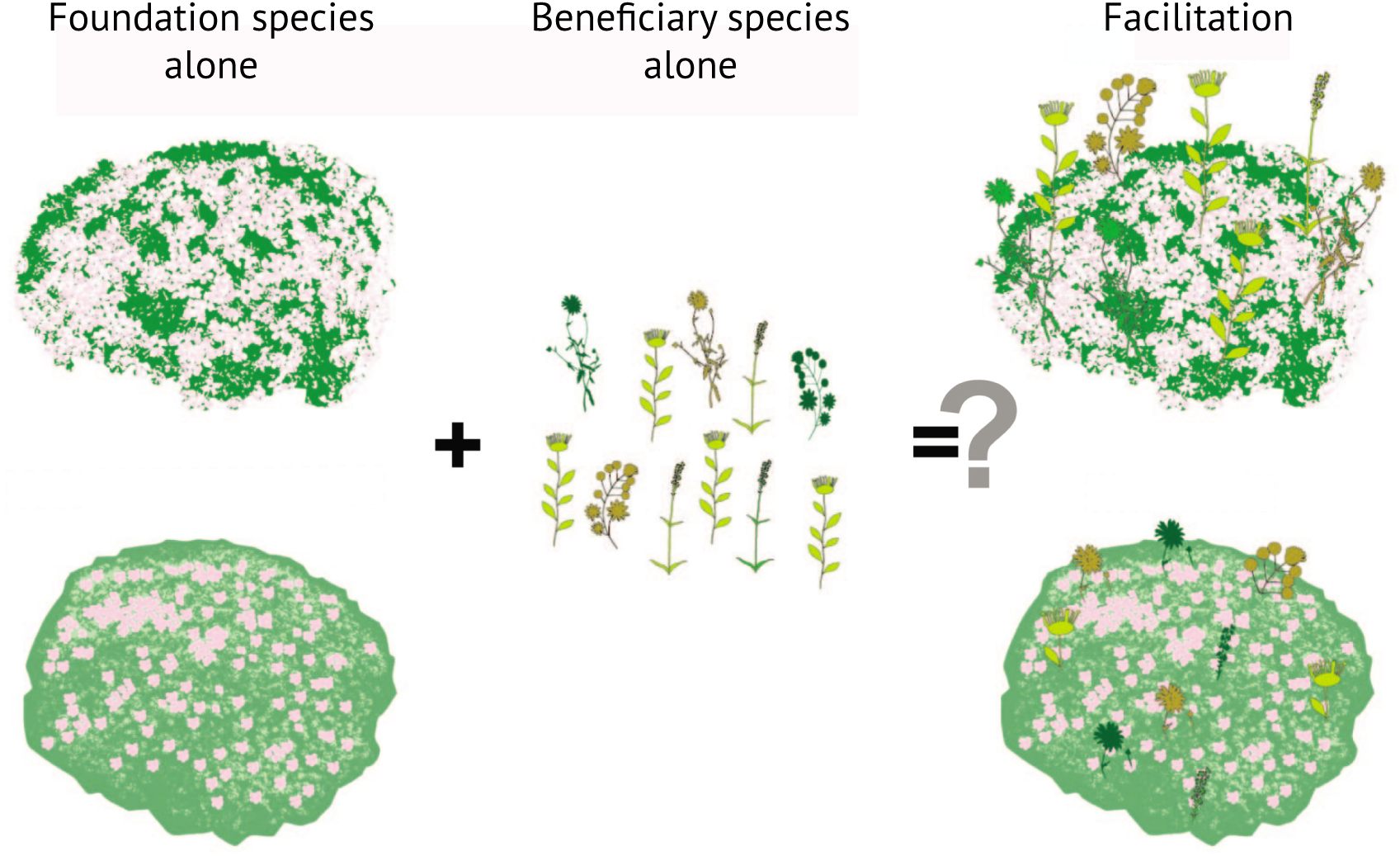
Overview of the experimental design to assess non-additive effects among plant species on pollinator networks. ‘Foundation species alone’: a cushion of the foundation species *Are-naria tetraquetra* spp. *amabilis* or *Hormathophylla spinosa* growing alone; ‘Beneficiary species alone’: non-cushion plant species growing alone; ‘Facilitation’: *Arenaria* or *Hormathophylla* and associated beneficiary species growing together.

## Results

### Pollinator diversity

We found that pollinator diversity significantly differed between the observed facilitation clusters and the expected additive sum depending on the identity of foundation species (F_1,52_ = 5.96, *p* = 0.0017, Figure 2, Tab. SI1). In particular, the facilitation clusters of Arenaria attracted a pollinator community that was *c.* 60 % more diverse than the additive expectation given by the simple sum of foundation and beneficiary species growing separately (q = 0.59, *p* = 0.0187). Differences were not significant for *Hormathophylla* (q = *-*0.32, *p* = 0.3600). These results suggest that in the case of facilitation by *Arenaria* net effects on pollinator diversity are synergistic rather than additive.

**Figure 2:**
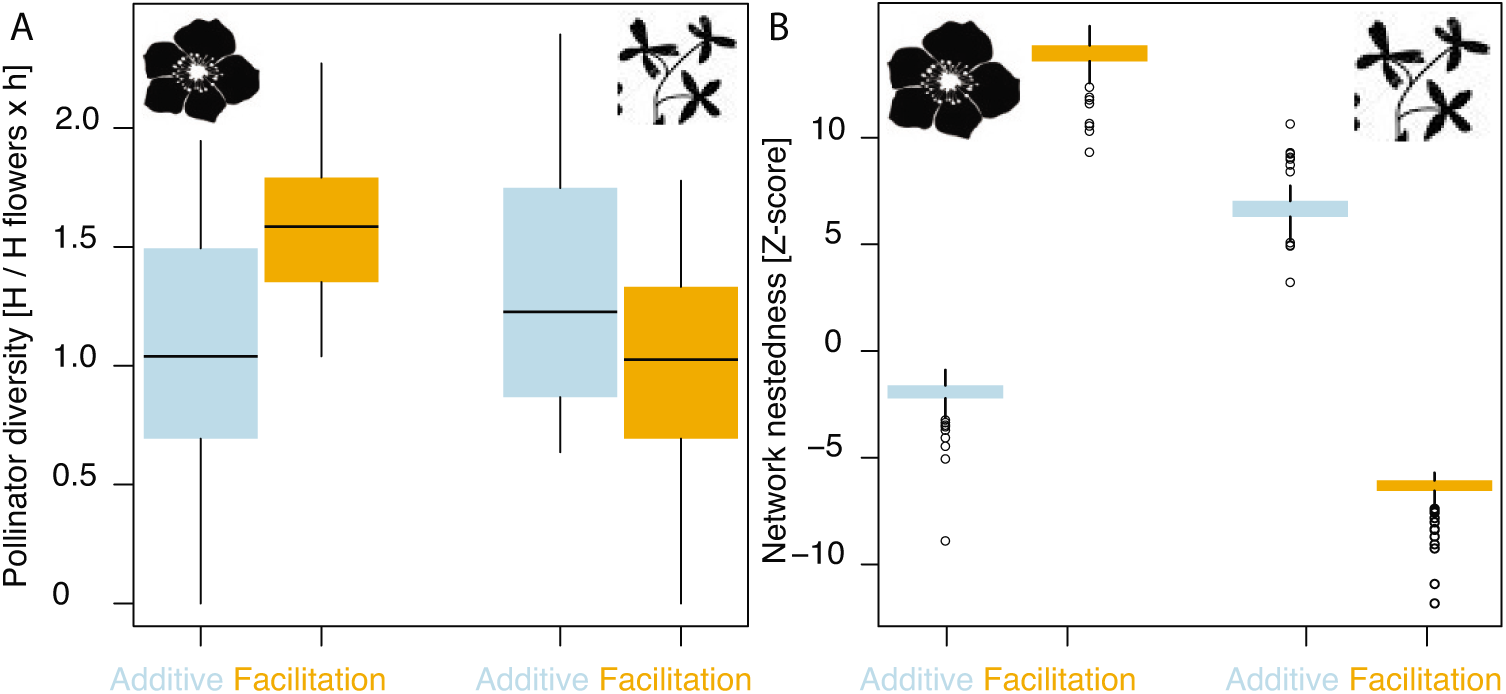
Differences between the expected additive community and the observed facilitation clusters in pollinator diversity and network architecture. (A) Pollinator Shannon diversity in expected networks assuming additive effects (light blue) and in observed facilitation networks (orange) of *Arenaria* (left) and *Hormathophylla* (right). Expected additive networks were obtained by summing the networks of foundation species alone and beneficiary species alone. (B) Nestedness of plant–pollinator networks. To compare nestedness among networks of different size, we calculated the relative nestedness (i.e., the Z-score, see Methods). In the box plots, horizontal bars show the median, the box the interquartile range and the vertical lines *±* 3 sd.

### Visitation rate

For each plant species, we assessed the potential beneficial or detrimental effects of growing in facilitation clusters for pollinator attraction (Fig. SI2). The pollinator visitation rate differed between treatments depending on the foundation species (F_1,176_ = 4.24, *p* = 0.0409) with average positive and negative effects in Arenaria and Hormathophylla, respectively (Fig. SI2, Tab. SI1). This indicates that chances of getting visited by pollinators varied when plant species grew in facilitation clusters or not, with contrasting consequences depending on foundation species. For instance, for *Jasione amethystina* and *Lotus corniculatus* spp. *glacialis* pollinator visitation rate increased and decreased under facilitation conditions in *Arenaria* and *Hormathoyphylla*, respectively.

### Network architecture

To analyze architectural changes between observed facilitation networks and expected additive networks we compared the relative nestedness (Figure 2). Relative nestedness significantly changed between facilitation and additive networks (F_1,396_ = 175.85, *p <* 0.0001) with differences between the two foundation species (interaction term: F_1,396_ = 21179.7, *p <* 0.0001, Tab. SI1). Specifically, *Arenaria* facilitation clusters showed a 16-fold increase in network nestedness in comparison with what would be expected assuming additive effects (q = 15.88, *p <* 0.0001). In contrast, *Hormathophylla* facilitation clusters were 13-fold less nested than the expected additive networks (q = *-*13.23, *p <* 0.0001). Taken together, these results demonstrate that observed facilitation clusters show also non-additive effects on pollination network architecture in contrasting ways depending on foundation species identity.

### Species turnover and interaction rewiring

In order to examine the potential mechanisms that might explain non-additive effects and their consequences for network robustness, we disentangled differences due to changes in species composition from differences due to interaction rewiring, i.e., the changes in interactions between plants and pollinators present in both additive and facilitation networks. Hence, we first quantified the network dissimilarity between expected and observed networks using the beta diversity of interactions (Poisot *et al.*, 2012). Network dissimilarity was 42.3 % for both foundation species (Fig. SI3). In *Arenaria*, 20.0 % of this difference was due to interaction rewiring and 22.3 % to species turnover. In *Hormathohylla*, 25.0 % was due to interaction rewiring and 17.3 % to species turnover. These results indicate that networks are different between treatments because they have both different species and because the species they share show different interactions.

### Interaction diversity

Having shown that changes in interactions contribute to differences between networks, we proceeded with examining networks composed only by species common to both additive and facilitation networks (Figure 3), in order to exclude differences due to species richness and composition. We found that species-level interaction diversity significantly differed between treatments and foundation species (F_1,57_ = 10.94, *p* = 0.0016, Figure 4A, Tab. SI1). Specifically, interaction diversity was higher in *Arenaria* clusters than expected by additive effects (q = 0.48, *p* = 0.0044) but as expected in *Hormathophylla* clusters (q = *-*0.17, *p* = 0.6144, Tab. SI2). These results indicate that plant facilitation increases the diversity of plant–pollinator interactions in the case of *Arenaria*, while the general effect was neutral in *Hormathophylla*. This went along with an overall increase in the generalization level of interactions within *Arenaria* clusters.

**Figure 3:**
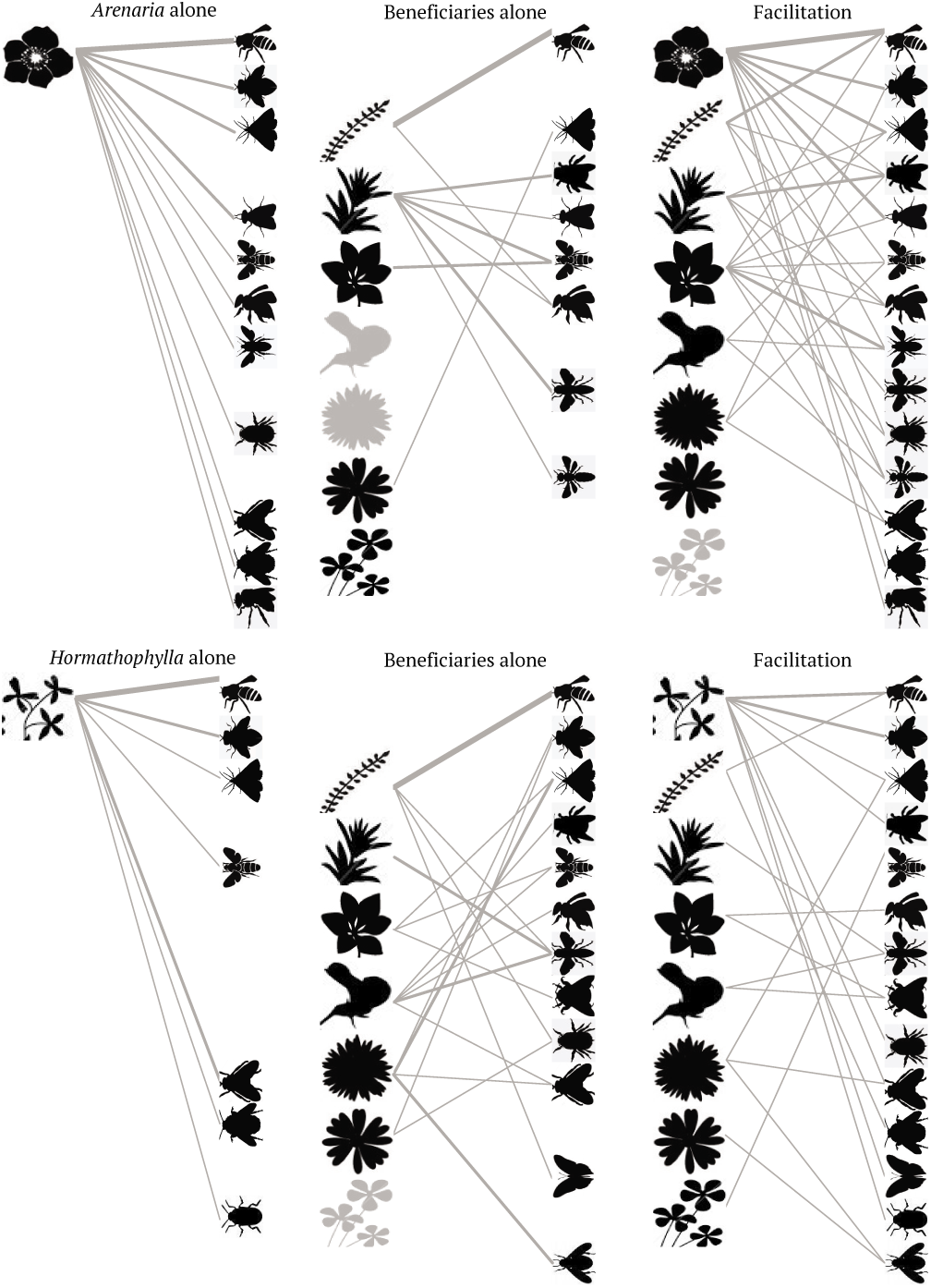
Plant–pollinator networks in experimental treatments composed only by pollinators common to both the observed facilitation and expected additive networks (i.e., the sum of foundation species alone and beneficiaries alone). The width of the links is proportional to interaction strength, measured as number of pollinators visiting a flower during one hour. Plants in black and without links were visited only in the network consisting of the entire pollinator species pool. Plants in gray were present but were not visited. For species names, see Fig. SI1. A visual inspection highlights the higher complexity of facilitation networks in *Arenaria* (above) and the lack thereof in *Hormathophylla* (below) compared to the expected sum of ?alone? networks.

**Figure 4:**
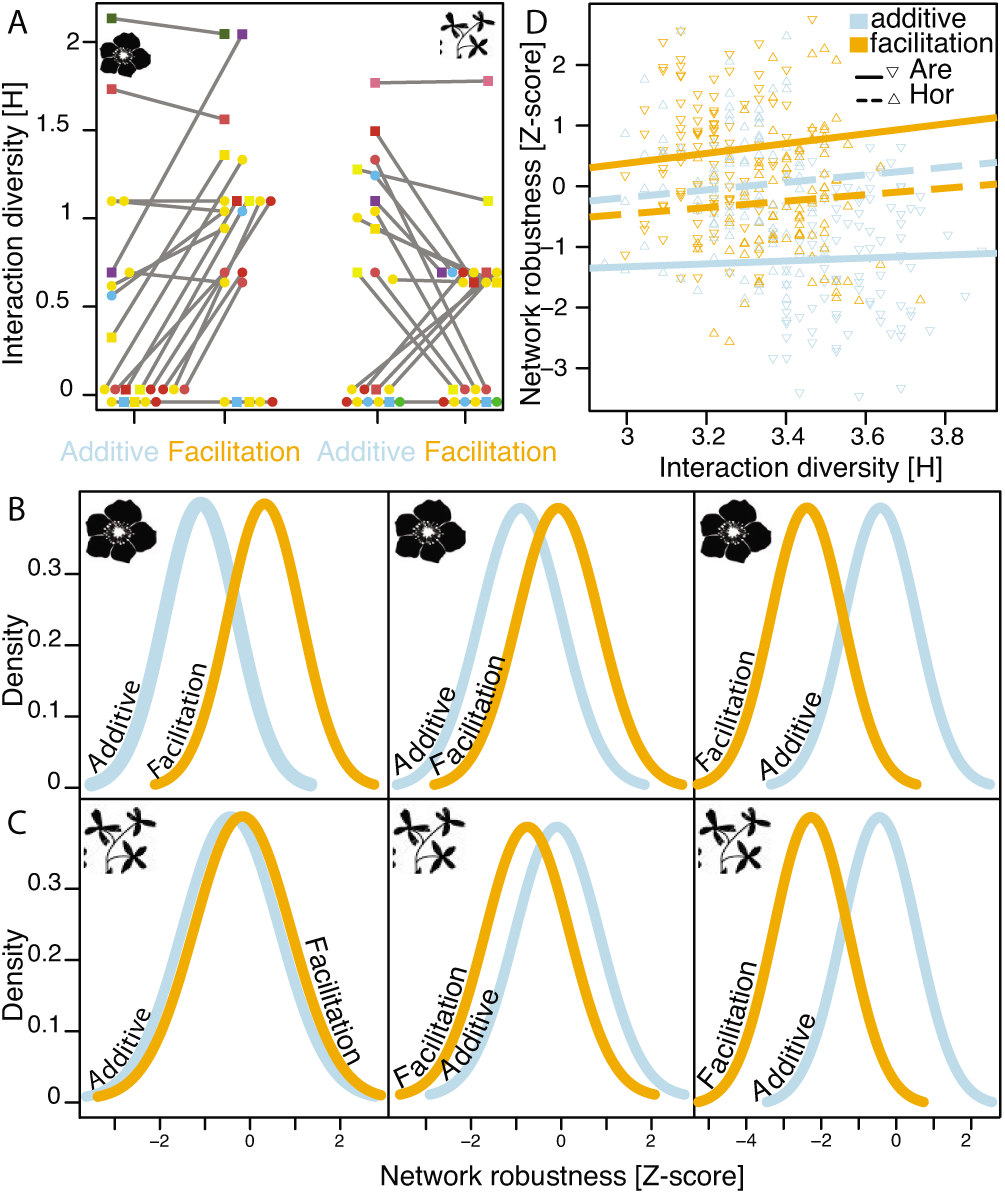
Properties of networks composed of species common between the additive and facilitation networks. (A) Interaction diversity of plant and pollinator species in expected additive networks and observed facilitation networks of *Arenaria* (left) and *Hormathophylla* (right). (B,C) Simulated network robustness against species loss (see Methods) of observed facilitation (orange) and expected additive (light blue) networks of *Arenaria* (B) and *Hormathophylla* (C) considering the scenarios of random extinctions (left), specialized-plants extinction (i.e., ad-dressing the pollinator community robustness; middle) and specialized-pollinators extinction (i.e., addressing the plant community robustness; right). (D) The strength of the relationship between network-level interaction diversity and network robustness varies between networks and foundation species depending on the scenario (Table SI3). Only the case of the random scenario is shown.

### Network robustness

Network robustness against species loss differed between treatments and foundation species with net effects varying depending on extinction scenarios (F_2,792_ = 33.67, *p <* 0.0001, Figure 4 B,C). In five cases out of six, the bottom-up effects of plant facilitation on pollination networks resulted in significantly different network robustness than what was expected from additive effects of foundation and beneficiary species (Tab. SI2). Specifically, in a random scenario the facilitation networks were c. two times more robust than expected for *Arenaria* (q = 1.75, *p <* 0.0001) but not for *Hormathophylla* (q = *-*0.23, *p* = 0.3636). In extinction scenarios that followed the relative abundance of interactions of plant and pollinator species (i.e., specialized scenario), respectively, pollinator and plant communities in *Arenaria* facilitation networks were 90 % more (q = 0.91, *p <* 0.0001) and two times less (q = *-*2.02, *p <* 0.0001) robust than additive networks, respectively, while in *Hormathophylla* facilitation networks were significantly less robust in both scenarios (q = *-*0.70, *p* = 0.0001; q = *-*1.83, *p <* 0.0001).

In addition, we found that the relationship between interaction diversity and network ro-bustness varied between treatments and foundation species (F_1,395_ = 57.82, *p <* 0.0001, Figure 4D; F_1,395_ = 15.93, *p <* 0.0001; F_1,395_ = 59.79, *p <* 0.0001 for random, specialized plants and specialized pollinators scenarios, respectively). Consequently, the diversity of interactions affected network robustness, regardless of species diversity, in different ways than assuming additive effects. Interestingly, the strength of this relationship varied between networks and foundation species depending on the extinction scenarios (Table SI3).

### The identity of foundation species and the directionality of non-additive effects

We suggest that the differences in the effects of *Arenaria* and *Hormathophylla* on pollinators may be due to the different position of the flowers of the associated beneficiary species within the canopy of the two foundation species (F_1,43_ = 30.05, *p <* 0.0001, Figure 5). In *Arenaria*, beneficiary species flower on top of the cushion canopy. Conversely, in *Hormathophylla*, the flowers of beneficiary species rarely reach the canopy and stay beneath. the flowers of beneficiary species rarely reach the canopy and stay beneath. This may result in non-additive effects with a synergistic outcome for *Arenaria* and antagonistic to neutral effects for *Hormathophylla*.

**Figure 5:**
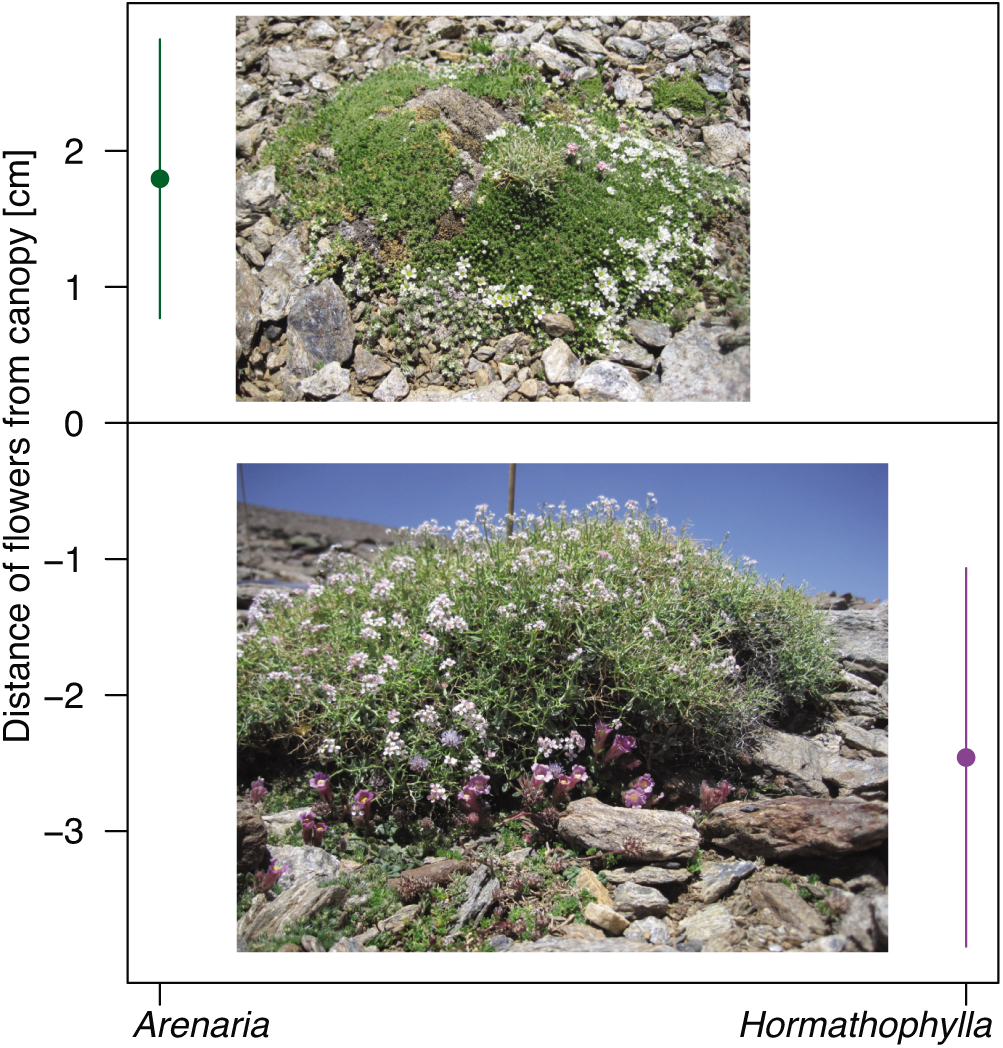
Distance of flowers of beneficiary species from the canopy of *Arenaria* (left and top) and *Hormathophylla* (right and bottom). Shown are 95 % CIs and pictures of the two foundation species with beneficiary species. In compact canopy of *Areanaria*, beneficiary species grow on top of it. Conversely, in the loose canopy of *Hormathophylla*, beneficiary species grow underneath.

## Discussion

We found that plant clustering through facilitation produced non-additive effects that scale up to pollination networks. Observed plant–pollinator interactions were different from expectations based on the sum of foundation and beneficiary species growing separately. This implies that plant–plant facilitation significantly affected plant–pollinator interactions, which in turn affected the architecture and robustness of plant–pollinator networks. The observed plant–pollinator interactions showed a certain degree of plasticity, that is variability in the identity of interacting partners. The existence of non-additive effects furthermore implies that interactions within a community are different from interactions outside a community context. In our specific case, the directionality of these effects, whether synergistic or antagonistic, depended on the identity of the foundation species, and in the case of robustness also on the extinction scenario.

These findings have implications for broader issues related to the nature of species interactions and the mechanisms regulating biodiversity in natural ecosystems. We argue that reductionist studies of species and their pairwise interactions in isolation from the community context might result in misleading conclusions.

First, our results support the hypothesis that positive plant–plant interactions can influence the assembly of the pollinator community as well as shape plant–pollinator networks. These results are in accordance with other studies showing the beneficial effects of foundation species on insect communities (Molina-Montenegro *et al.*, 2008; Reid & Lortie, 2012; Ruttan *et al.*, 2016). Overall, we found that positive plant interactions can scale up to modify plant–pollinator network architecture and robustness thanks to synergistic effects of plant aggregation and pollinator adaptive responses. Indeed, plant clusters created by facilitation increased the diversity of pollinator species and resulted in more nested plant–pollinator networks in one of two model systems. Furthermore, higher nestedness in the observed facilitation network with *Arenaria* may result in a reduction of competition among plants for pollinators (Bastolla *et al.*, 2009).

Second, we found that the diversity of plant–pollinator interactions affected network robustness regardless of the diversity of species. These results demonstrate that the diversity of species interactions is also relevant for ecosystem stability, an effect often attributed to the diversity of species *per se* (Hooper *et al.*, 2005). High interaction diversity can contribute to network robustness by creating redundancy of links among plants and pollinators. Such redundancy can be achieved on the one hand by higher species diversity, but on the other hand also by increased interaction plasticity of the interacting partners. Indeed, in case of *Arenaria*, we observed not only higher species diversity but also a shift in interaction plasticity, with pollinator interactions in facilitation clusters becoming more generalist than expected. Consequently, facilitation clusters can produce a magnet effect (Laverty, 1992; Molina-Montenegro *et al.*, 2008) that results in increased plant–pollinator interactions.

A third aspect of our results is that the negative effects of plant–plant competition for resources in plant clusters may be counterbalanced by the positive effects of plant–plant facilitation for pollination networks, finally contributing to the overall facilitative effect of foundation species on plant species richness and density (Schöb *et al.*, 2012). On the one hand, the higher density in plant clusters may increase competition among plants for resources (Grace & Tilman, 1990; Levine & Rees, 2002; Harpole & Tilman, 2006). On the other hand, we demonstrate that plant clustering can be beneficial for pollination (see also Laverty, 1992; Feldman *et al.*, 2004; Mesgaran *et al.*, 2017), hence potentially increasing sexual reproduction of plants (Schöb *et al.*, 2014).

The coexistence of the competing plant species can be explained by the ‘cluster effect’ (a socio-economical concept *sensu* Porter 1998; 2008) where, in our case, benefits in the pollination service overcome the negative impacts of plant competition. In other words, our study demonstrates that species clusters, such as those created by foundation species, cannot be explained by simple pairwise interactions but request an understanding of the higher order interactions (Levine *et al.*, 2017), such as the role of pollinators in interactions among plants.

## Material and methods

### Experimental setting

A selective removal experiment was performed in the Sierra Nevada Mountains (Loma del Mulhacén, Spain) during July 2015. The study site is located at 3200 m a.s.l. (Lat 37.041417N, Long-003.306400W), characterised by a patchy alpine vegetation dominated by the cushion-forming species *Arenaria tetraquetra* ssp. *amabilis* (Bory) H. Lindb. Fil. (Caryophyllaceae) and *Hormathophylla spinosa* (L.) Kupfer (Brassicaceae). These foundation species provide positive facilitative effects on other beneficiary plant species through the improvement of their physiological status (Schöb *et al.*, 2012) and reproductive output (Schöb *et al.*, 2014). These facilitation mechanisms are due to the decrease of stress followed by the increase of soil water content and organic matter in foundation species compared to bare ground (Schöb *et al.*, 2013a,b).

Our null hypothesis is that facilitation influences pollination. Our null expectation is that the pollinator community in a facilitative system is the sum of the components of the facilitative system (i.e., foundation species and beneficiary species). Therefore, we considered the naturally-occurring facilitation clusters with foundation species and beneficiary species as control. In the removal treatment we either removed the foundation species (to have the beneficiary species growing alone) or we removed the beneficiary species (to have the foundation species growing alone). Each treatment consisted of a standard plot size of 20 x 20 cm. Distance among plots within block ranged between 0.5 m and 1 m. We followed a randomised block design, where each block was composed by foundation species and beneficiary species growing separately and foundation species and beneficiary species growing together replicated over the two foundation species (Figure 1). In total, 14 blocks were established within a relatively homogeneous area of about 1 ha, resulting in 84 plots in total. Plant species composition is the same overall and was kept as similar as possible between treatments of the same block (Tab. SI4).

Plant–pollinator interactions were observed during the entire flowering season of July 2015. Thanks to an exceptionally dry spring and a warm summer, plants completed their flowering phase within three weeks during July. Hence, we were able to cover the complete flowering time for most of the species at our study site. Each plot was sampled during a standardised time span of 20 min a day. The three plots belonging to the same block were sampled together, in order to eliminate within block variability due to sampling weather conditions. Every day 14 sampling rounds were carried out between 10 am and 5.30 pm (blocks randomly sampled). Each block was sampled between 6 and 9 times, resulting in 204 sampling rounds in total (Tab. SI5). All flower-visiting insects of each flower (plant species) in each plot were sampled using a sweep net or an entomological aspirator. Thus, pollinator specimens were attributed to a specific plant species within each plot (Tab. SI6). Due to conservation issues related to Sierra Nevada National Park legislation and also ethical issues, we limited the collection of bees, bumblebees, hoverflies and butterflies to those necessary for species identification. Insects were identified at the species level whenever possible, otherwise to genus or family (Tab. SI6). As not all the flower-visiting insects are actual pollinators, we excluded from the analysis all the not-pollinator species on the basis of expert knowledge (Tab. SI6). Insect specimens are stored at the ETH insect collection and at our private collections.

### Network analysis

To assess wether the observed community is different from the sum of its single components, we compared pollinator communities visiting the observed facilitation clusters with foundation and beneficiary species (i.e., control, ‘facilitation’ treatment in Figure 1) with the expected additive pollinator communities calculated as the sum of the species growing alone. We highlight that the comparison of the observed facilitation clusters to the sum of its components is a more conservative approach than that of a mean of the components as we have double the area for the expected community. However, we believe that this approach is not only more conservative but also more accurate because we keep the number of plants and flowers similar among treatments.

Pollinator diversity was calculated with the Shannon index (Oksanen *et al.*, 2017) at the plot level. As pollinator abundance responds to flower density (Losapio *et al.*, 2016), visitation rate of each plant species was calculated by standardising the number of pollinators by the number of flowers and sampling hours at the plot level. Then, we considered the pollination networks at the treatment level. Network architecture was calculated according to the measure of nestedness by Bastolla *et al.* (Bastolla *et al.*, 2009). We chose this metric instead of NODF because the latter does not accurately consider the contribution of different species with the same degree to network nestedness, which in our case is fundamental given the long tail of pollinators with few visits per plant species. This nestedness was calculated as 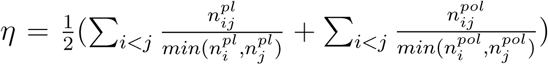 where *n_ij_*is the number of interactions *n* between two plant (pl) or two pollinator (*pol*) species *i*–*j* and min(*n*_*i*_*, n*_*j*_) is the smaller of the two values. To estimate the significance of each observed network, we compared the observed index with the distribution (95 % confidence interval) of 100 random networks (Tab. SI4). Random networks were built according to a probabilistic null model (Bascompte *et al.*, 2003), which does a relative good job in minimising simultaneously Type I and Type II errors (Rodríguez-Gironés & Santamaría, 2006) and it is most biologically meaningful in terms of species generalisation (i.e., node degree). This null model builts networks from a template of interaction probabilities, such that in an adjacency matrix *A* = *R × C* with *R* and *C* rows and columns, respectively, the probability that a cell *a*_*ij*_ has a link is 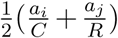 where *a*_*i*_ and *a*_*j*_ is the number of links in row *i* and column *j*, respectively. Only random networks with *R* and *C* equal to the empirical networks were considered. To compare nestedness between facilitation and additive networks, we controlled for the variation in matrix size (*R, C*) by calculating the deviance of the empirical nestedness with the random expectation given by the 100 replicates of the probabilistic null model as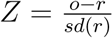, where *o* and *r* are the values of epirical and random networks, respectively, weighted by the standard deviation *sd* of random networks.

Second, we focused on the networks composed by the species common to both treatments. We first decomposed network dissimilarity (Fig. SI3) into species turnover and interaction rewiring components using the *β*-diversity of interactions approach (Poisot *et al.*, 2012). Thus, to exclude differences between networks due to changes in species composition we built networks considering only shared species between treatments (Figure 3). To estimate interaction rewiring, we then calculated both the species-level and network-level diversity of interactions using the Shannon index (Oksanen *et al.*, 2017). To assess the functional consequences of such interaction rewiring we computed the robustness of networks against secondary extinctions. In absence of biologically-informed criteria of species susceptibility to environmental perturbations (Losapio & Schöb, 2017), we proceeded considering a scenario of random extinction of species, a scenario of extinction in order of most specialised plants and a third of most specialised pollinators (Solé & Montoya, 2001; Albert & Barabasi, 2002; Dunne *et al.*, 2002). Robustness was calculated as the area under the secondary extinction curve generated by sequential removal of plant and pollinator species in random order (Dormann *et al.*, 2009). Each series was replicated 100 times. Each empirical value was compared to a distribution (95 % confidence interval) of 100 random networks (Z-score).

### Statistical analysis

Regression models were used to assess the effect of treatment (additive vs. facilitation), foundation species (i.e., *Arenaria* or *Hormathophylla*) (predictors) and their interactions on pollinator diversity, relative nestedness (responses, two different models). To assess changes in pollinator visitation rate and interaction diversity (responses) a mixed-effect model approach was used fitting species identity as random term and treatment and foundation species as fixed effects. To assess differences in network robustness (response) a mixed-effect model was fitted using extinction scenario, treatment, foundation species and their interactions as predictors and the random network as error term. The effect of network-level interaction diversity (predictor) on relative robustness (response) was tested using a regression model with the interaction terms interaction diversity–treatment and interaction diversity–foundation species for each scenario separately. To assess the significance of specific contrasts, Tukey HSD post-hoc tests and comparisons among least-squares means (with Tukey correction) were performed on each statistical model. All data analyses were performed in R version 3.3.3 (R Core Team, 2017).

## Acknowledgements

This study was supported by the Swiss National Science Foundation awarded to CS (PZ00P3 148261). JB is supported by the European Research Council through an Advanced Grant. We thank L. Dutoit for helping with data collection, M. Furler for drawing Figure 1 and the Sierra Nevada National Park for providing sampling permissions. Thanks to Hannes Baur, Andreas Muüller, Martin Schwarz for helping with further species identification.

## Data accessibility

All original data used in this study will be available from the Dryad Digital Repository. **R scripts** will be uploaded as online supporting information.

## Supporting Information

Additional supporting information may be found in the online version of this article.

## Author contributions

G.L. and C.S. designed and performed the experiment; G.L., M.A.F., J.B., B.S., R.M. and C.S. discussed methodological issues and results; R.N., H.B., L.C., P.C., C.G., J.P.H., S.K., A.M., J.O., A.C.P., P.R., J.S., M.S. and D.S. contributed to species identification; G.L. analysed the data with input from M.A.F. and J.B.; G.L. wrote the first draft of the paper and all authors contributed to the subsequent revision.

The authors declare no conflict of interest.

## References

Albert, R. & Barabasi, A. L. (2002). Statistical mechanics of complex networks. Reviews of Modern Physics, 74, 47–97.

Bascompte, J. & Jordano, P. (2014). Mutualistic Networks. Princeton University Press, Princeton, New Jersey, USA.

Bascompte, J., Jordano, P., Melian, C. J. & Olesen, J. M. (2003). The nested assembly of plant-animal mutualistic networks. Proceedings of the National Academy of Sciences of the United States of America, 100, 9383–9387.

Bastolla, U., Fortuna, M. A., Pascual-Garcia, A., Ferrera, A., Luque, B. & Bascompte, J. (2009). The architecture of mutualistic networks minimizes competition and increases biodiversity. Nature, 458, 1018–1020.

Bruno, J. F., Stachowicz, J. J. & Bertness, M. D. (2003). Inclusion of facilitation into ecological theory. Trends in Ecology & Evolution, 18, 119–125.

Callaway, R. M. (2007). Positive Interactions and Interdependence in Plant Communities. Springer.

Cavieres, L. a., Brooker, R. W., Butterfield, B. J., Cook, B. J., Kikvidze, Z., Lortie, C. J., Michalet, R., Pugnaire, F. I., Schöb, C., Xiao, S., Anthelme, F., Björk, R. G., Dickinson, K. J. M., Cranston, B. H., Gavilán, R., Gutiérrez-Girón, A., Kanka, R., Maalouf, J. P., Mark, A. F., Noroozi, J., Parajuli, R., Phoenix, G. K., Reid, A. M., Ridenour, W. M., Rixen, C., Wipf, S., Zhao, L., Escudero, A., Zaitchik, B. F., Lingua, E., Aschehoug, E. T. & Callaway, R. M. (2014). Facilitative plant interactions and climate simultaneously drive alpine plant diversity. Ecology Letters, 17, 193–202.

Dormann, C. F., Frund, J., Bluthgen, N. & Gruber, B. (2009). Indices, Graphs and Null Models: Analyzing Bipartite Ecological Networks. The Open Ecology Journal, 2, 7–24.

Duchene, O., Vian, J.-F. & Celette, F. (2017). Intercropping with legume for agroecological cropping systems: Complementarity and facilitation processes and the importance of soil microorganisms. a review. Agriculture, Ecosystems & Environment, 240, 148–161.

Dunne, J. A., Williams, R. J. & Martinez, N. D. (2002). Network structure and biodiversity loss in food webs: Robustness increases with connectance. Ecology Letters, 5, 558–567.

Ellison, A. M., Bank, M. S., Clinton, B. D., Colburn, E. a., Elliott, K., Ford, C. R., Foster, D. R., Kloeppel, B. D., Knoepp, J. D., Lovett, G. M., Mohan, J., Orwig, D. a., Rodenhouse, N. L., Sobczak, W. V., Stinson, K. a., Stone, J. K., Swan, C. M., Thompson, J., VonHolle, B. & Webster, J. R. (2005). Loss of foundation species: Consequences for the structure and dynamics of forested ecosystems. Frontiers in Ecology and the Environment, 3, 479–486.

Feldman, T. S., Morris, W. F. & Wilson, W. G. (2004). When can two plant species facilitate each other’s pollination? Oikos, 105, 197–207.

Grace, J. B. & Tilman, D. (1990). Perspectives on Plant Competition. Academic Press, 24–28 Oval Road, London NW1 7DX.

Hacker, S. D. & Gaines, S. D. (1997). Some implications of direct positive interactions for community species diversity. Ecology, 78, 1990–2003.

Harpole, W. S. & Tilman, D. (2006). Non-neutral patterns of species abundance in grassland communities. Ecology Letters, 9, 15–23.

Hector, A., Schmid, B., Beierkuhnlein, C., Caldeira, M. C., Diemer, M., Dimitrakopoulos, P. G., Finn, J. A., Freitas, H., Giller, P. S., Good, J., Harris, R., Hogberg, P., Huss-Danell, K., Joshi, J., Jumpponen, A., Körner, C., Leadley, P. W., Loreau, M., Minns, A., Mulder, C. P. H., O’Donovan, G., Otway, S. J., Pereira, J. S., Prinz, A., Read, D. J., Scherer-Lorenzen, M., Schulze, E. D., Siamantziouras, A. S. D., Spehn, E. M., Terry, A. C., Troumbis, A. Y., Woodward, F. I., Yachi, S. & Lawton, J. H. (1999). Plant diversity and productivity experiments in european grasslands. Science, 286, 1123.

Hooper, D. U., Chapin, F. S., Ewel, J. J., Hector, A., Inchausti, P., Lavorel, S., Lawton, J. H., Lodge, D. M., Loreau, M., Naeem, S., Schmid, B., Setala, H., Symstad, A. J., Vandermeer, J. & Wardle, D. A. (2005). Effects of biodiversity on ecosystem functioning: a consensus of current knowledge. Ecological Monographs, 75, 3–35.

Laverty, T. M. (1992). Plant interactions for pollinator visits: a test of the magnet species effect. Oecologia, 89, 502–508.

Levine, J. M., Bascompte, J., Adler, P. B. & Allesina, S. (2017). Beyond pairwise mechanisms of species coexistence in complex communities. Nature, 546, 56–64.

Levine, J. M. & Rees, M. (2002). Coexistence and relative abundance in annual plant assemblages: the roles of competition and colonization. The American Naturalist, 160, 452–67.

Losapio, G., Gobbi, M., Marano, G., Avesani, D., Boracchi, P., Compostella, C., Pavesi, M., Schöb, C., Seppi, R., Sommaggio, D., Zanetti, A. & Caccianiga, M. (2016). Feedback effects between plant and flower-visiting insect communities along a primary succession gradient. Arthropod-Plant Interactions, 10, 485–495.

Losapio, G. & Schöb, C. (2017). Resistance of plant–plant networks to biodiversity loss and secondary extinctions following simulated environmental changes. Functional Ecology, 31, 1145–1152.

MacArthur, R. & Levins, R. (1967). The limiting similarity, convergence, and divergence of coexisting species. The American Naturalist, 101, 377–385.

McIntire, E. J. B. & Fajardo, A. (2014). Facilitation as a ubiquitous driver of biodiversity. New Phytologist, 201, 403–416.

Meron, E. (2012). Pattern-formation approach to modelling spatially extended ecosystems. Ecological Modelling, 234, 70–82.

Mesgaran, M. B., Bouhours, J., Lewis, M. A. & Cousens, R. D. (2017). How to be a good neighbour: Facilitation and competition between two co-flowering species. Journal of Theoretical Biology, 422, 72–83.

Michalet, R., Brooker, R. W., Cavieres, L. a., Kikvidze, Z., Lortie, C. J., Pugnaire, F. I., Valiente-Banuet, A. & Callaway, R. M. (2006). Do biotic interactions shape both sides of the humped-back model of species richness in plant communities? Ecology Letters, 9, 767–773.

Molina-Montenegro, M. A., Badano, E. I. & Cavieres, L. A. (2008). Positive interactions among plant species for pollinator service: assessing the ‘magnet species’ concept with invasive species. Oikos, 117, 1833–1839.

Oksanen, J., Guillaume Blanchet, F., Friendly, M., Kindt, Legendre P., McGlinn, D., Minchin, P. R., O’Hara, R. B., Simpson, G. L., Solymos, P., Stevens, M. H. H., Szoecs, E. & Wagner, H. (2017). vegan: Community ecology package. R package version 2.4-2.

Poisot, T., Canard, E., Mouillot, D., Mouquet, N. & Gravel, D. (2012). The dissimilarity of species interaction networks. Ecology Letters, 15, 1353–1361.

Porter, M. E. (1998). Clusters and the new economics of competition. Harvard Business Review, 98609, 77–90.

Porter, M. E. (2008). On Competition. Harvard Business School Publishing Corporation.

R Core Team (2017). R: A Language and Environment for Statistical Computing. URL http://www.r-project.org/.

Reid, A. M. & Lortie, C. J. (2012). Cushion plants are foundation species with positive effects extending to higher trophic levels. Ecosphere, 3.

Rodríguez-Gironés, M. A. & Santamaria, L. (2006). A new algorithm to calculate the nestedness temperature of presence–absence matrices. Journal of Biogeography, 33, 924–935.

Ruttan, A., Filazzola, A. & Lortie, C. J. (2016). Shrub-annual facilitation complexes mediate insect community structure in arid environments. Journal of Arid Environments, 134, 1–9.

Schöb, C., Armas, C., Guler, M., Prieto, I. & Pugnaire, F. I. (2013a). Variability in functional traits mediates plant interactions along stress gradients. Journal of Ecology, 101, 753–762.

Schöb, C., Armas, C. & Pugnaire, F. I. (2013b). Direct and indirect interactions co-determine species composition in nurse plant systems. Oikos, 122, 1371–1379.

Schöb, C., Butterfield, B. J. & Pugnaire, F. I. (2012). Foundation species influence trait-based community assembly. New Phytologist, 196, 824–834.

Schöb, C., Kerle, S., Karley, A. J., Morcillo, L., Pakeman, R. J., Newton, A. C. & Brooker,R. W. (2015). Intraspecific genetic diversity and composition modify species-level diversity–productivity relationships. New Phytologist, 205, 720–730.

Schöb, C., Prieto, I., Armas, C. & Pugnaire, F. I. (2014). Consequences of facilitation: one plant’s benefit is another plant’s cost. Functional Ecology, 28, 500–508.

Sieber, Y., Holderegger Rolf, R., Waser, N. M., Thomas, V. F. D., Braun, S., Erhardt, A., Reyer, H. U. & Wirth, L. R. (2011). Do alpine plants facilitate each other’s pollination? Experiments at a small spatial scale. Acta Oecologica, 37, 369–374.

Solé, R. V. & Montoya, J. M. (2001). Complexity and fragility in ecological networks. Proceedings of the Royal Society B: Biological Sciences, 268, 2039–2045.

Valiente-Banuet, A., Rumebe, A. V., Verdú, M. & Callaway, R. M. (2006). Modern quaternary plant lineages promote diversity through facilitation of ancient tertiary lineages. Proceedings of the National Academy of Sciences, 103, 16812–16817.

Verdú, M. & Valiente-Banuet, A. (2008). The nested assembly of plant facilitation networks prevents species extinctions. The American Naturalist, 172, 751–760.

